# Dynamics of isoflurane-induced vasodilation and blood flow of cerebral vasculature revealed by multi-exposure speckle imaging

**DOI:** 10.1101/2020.06.26.174227

**Authors:** Colin T. Sullender, Lisa M. Richards, Fei He, Lan Luan, Andrew K. Dunn

**Affiliations:** Department of Biomedical Engineering, University of Texas at Austin, 107 W. Dean Keeton Street Stop C0800, Austin, TX, 78712, United States; Department of Electrical and Computer Engineering, Rice University, 6100 Main Street, Houston, TX, 77005, United States; Department of Bioengineering, Rice University, 6100 Main Street, Houston, TX, 77005, United States

**Keywords:** multi-exposure speckle imaging, laser speckle contrast imaging, awake imaging, anesthesia, hemodynamics, cerebral blood flow

## Abstract

**Background:** Anesthetized animal models are used extensively during neurophysiological and behavioral studies despite systemic effects from anesthesia that undermine both accurate interpretation and translation to awake human physiology. The majority of work examining the impact of anesthesia on cerebral blood flow (CBF) has been restricted to before and after measurements with limited spatial resolution.

**New Method:** We used multi-exposure speckle imaging (MESI), an advanced form of laser speckle contrast imaging (LSCI), to characterize the dynamics of isoflurane anesthesia induction on cerebral vasculature and blood flow in the mouse brain.

**Results:** The large anatomical changes caused by isoflurane are depicted with wide-field imagery and video highlighting the induction of general anesthesia. Within minutes of exposure, both vessel diameter and blood flow increased drastically compared to the awake state and remained elevated for the duration of imaging. An examination of the dynamics of anesthesia induction reveals that blood flow increased faster in arteries than in veins or parenchyma regions.

**Comparison with Existing Methods:** MESI offers robust hemodynamic measurements across large fields-of-view and high temporal resolutions sufficient for continuous visualization of cerebrovascular events featuring major changes in blood flow.

**Conclusion:** The large alterations caused by isoflurane anesthesia to the cortical vasculature and CBF are readily characterized using MESI. These changes are unrepresentative of normal physiology and provide further evidence that neuroscience experiments would benefit from transitioning to un-anesthetized awake animal models.

## 1. Introduction

The use of general anesthesia during neuroimaging is ubiquitous across many animal models despite systemic effects on neurophysiological state and cardiovascular function [1]. Volatile inhalation anesthetics, such as the halogenated ether isoflurane [2], are commonly utilized to immobilize animals during imaging while allowing for fine control over the depth of anesthesia and consciousness. However, isoflurane has been shown to reduce neuronal activity [3] and functional connectivity [4], suppress the magnitude and speed of neurovascular coupling [5, 6, 7], and induce significant vasodilation [8, 9]. Isoflurane also conveys potential neuroprotective effects that reduce and delay the severity of cerebral ischemia [10, 11, 12, 13, 14]. These effects can mask the benefits of prospective neuroprotective therapeutics or interventions and confound the outcomes of long-term studies [15, 16]. For these reasons and more [17], there is a growing effort within the neuroscience community to transition to un-anesthetized awake animal models in order to more accurately interpret neurophysiological and behavioral experiments and more readily translate findings to awake human physiology.

There have been numerous imaging modalities used to examine the effects of isoflurane on cerebral hemodynamics. Functional magnetic resonance imaging (fMRI), laser Doppler flowmetry, and intrinsic signal optical imaging have all established that isoflurane increases basal cerebral blood flow (CBF) while attenuating and delaying the hemodynamic response to local neural activity via neurovascular coupling [18, 3, 6, 7]. Two-photon phosphorescence lifetime microscopy showed that tissue oxygenation is twice as high under isoflurane anesthesia compared to the awake state and exhibited large layer-specific differences within the cortical vasculature [19]. Calcium signal imaging following ischemic stroke found that the magnitude of spreading depolarizations were significantly smaller in awake mice [20] while laser speckle contrast imaging (LSCI) of photothrombotic stroke observed larger infarct sizes in awake rats compared to their isoflurane-anesthetized counterparts [14]. To our knowledge, no prior studies have directly imaged the induction of general anesthesia and its acute effects upon the cerebral vasculature.

In this paper, we present multi-exposure speckle imaging (MESI) of cerebral blood flow in awake mice during the induction of general anesthesia with isoflurane. The MESI technique [21] provides a more robust estimate of large changes in flow compared to traditional single-exposure LSCI [22] and enables the chronic tracking of CBF [23]. We examine the dynamics of the large anatomical and physiological changes to cortical vasculature in response to isoflurane with wide-field imagery across multiple imaging sessions and animals. Because the anesthetized state is representative of an abnormal physiology, we argue that awake animal models should be utilized whenever possible for neuroscience studies.

## 2. Materials and Methods

### 2.1. Multi-Exposure Speckle Imaging (MESI)

A schematic of the imaging system is presented in Fig. 1a. MESI was performed using a 685 nm laser diode (50 mW, HL6750MG, Thorlabs, Inc.) intensity modulated with an acousto-optic modulator (AOM, 3100-125, Gooch & Housego, Ltd.) and relayed to illuminate the craniotomy at an oblique angle. The scattered light was imaged by a CMOS camera (acA1920-155um, Basler AG) with 2 × magnification and cropped to a field-of-view of 3.6 × 3.0 mm. Camera exposures were temporally synchronized with the modulated laser pulses. Fifteen camera exposures ranging between 50 *µ*s and 80 ms [21, 24, 23] were recorded for each complete MESI frame in order to broadly sample the speckle dynamics of the specimen [25], resulting in an effective acquisition rate of ∼2.5 MESI frames-per-second. The total amount of light used to capture each exposure was held constant with the AOM in order to minimize the effects of shot noise [21]. The acquisition was controlled using custom software written in C++ along with a multifunction I/O device (USB-6363, National Instruments Corp.) for the generation of camera exposure trigger signals and AOM modulation voltages [26].

**Figure 1:**
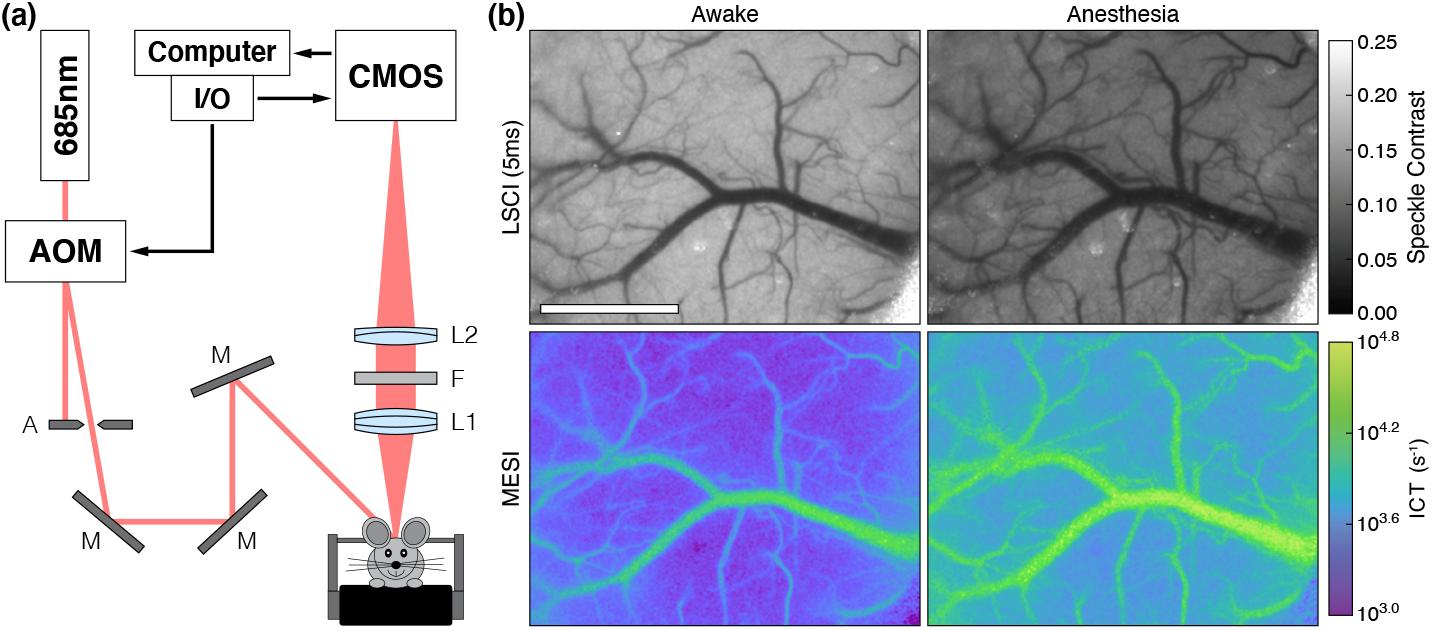
(a) MESI system schematic with 2× magnification: A (Standard Iris, ID15, Thor-labs, Inc.), M (Ø1” Broadband Dielectric Mirror, BB1-E02, Thorlabs, Inc.), L1 (*f* = 50 mm, Steinheil Triplet Achromatic, 67-422, Edmund Optics, Inc.), F (685 ± 20 nm bandpass filter, S685/40m, Chroma Technology Corp.), and L2 (*f* = 100 mm, Achromatic Doublet, AC254- 100-A, Thorlabs, Inc.). (b) Single exposure LSCI (5 ms) and MESI during the awake and anesthetized states in Subject 1 (Scale bar = 1 mm). See Video 1 (MPEG-4, 29.8 MB) for the transition between each state during the induction of general anesthesia.

The 15 raw intensity images were converted to speckle contrast images (*K* = *σ*_*s*_ / ⟨ *I* ⟩) with a 7 × 7-pixel sliding window and used to calculate an estimate of the speckle correlation time (*τ*_*c*_) at each pixel with the multi-exposure speckle visibility expression [21]:

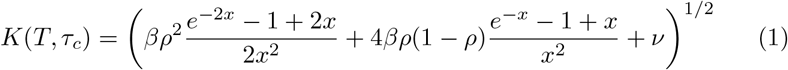

where *T* is the camera exposure time, *x* = *T/τ*_*c*_, *ρ* is the fraction of light that is dynamically scattered, *β* is a normalization factor that accounts for speckle sampling, and *ν* represents exposure-independent instrument noise and nonergodic variances. Eq. (1) was fitted with the Levenberg-Marquardt nonlinear least squares algorithm [27] using a custom program written in C.

Because *τ*_*c*_ is inversely related to the speed of the moving scatterers [28, 29], the inverse correlation time (*ICT* = 1*/τ*_*c*_) is frequently used as a metric for quantifying blood flow in vasculature and perfusion in parenchyma [30, 31, 26]. Recent work to improve the quantitative accuracy of MESI flow measurements in vasculature takes into account the presence of multiple dynamic scattering events instead of assuming only a single dynamic scattering event per photon [32]. This is achieved by scaling the fitted *ICT* value by the diameter of the vessel in order to obtain an estimate of “vascular flux” that better accounts for variations in vascular volume sampling [32, 33]. This paper uses the vascular flux metric (arbitrary units) for all vessel measurements and *ICT* (s^−1^) for all parenchyma perfusion measurements.

### 2.2. Animal Preparation

Mice (*n* = 4, C57BL/6J, male, 4-6 months, Charles River Laboratories, Inc.) were anesthetized with medical O_2_ vaporized isoflurane (3% for induction, 1-2% for maintenance, 0.5 L/min) via nose-cone inhalation. Body temperature was maintained at 37 ^◦^C with a feedback heating pad (DC Temperature Controller, Future Health Concepts, Inc.). Arterial oxygen saturation, heart rate, and breathing rate were monitored via pulse oximetry (MouseSTAT, Kent Scientific Corp.). Mice were placed supine in a stereotaxic frame (Kent Scientific Corp.) and subcutaneously administered carprofen (5 mg/kg) and dexamethasone (2 mg/kg) to reduce inflammation during the craniotomy procedure. The scalp was shaved and resected to expose the skull between the bregma and lambda cranial coordinates. A circular region (4 mm diameter) of the skull including the dura mater over the somatosensory cortex was removed with a dental drill (Ideal Microdrill, 0.8 mm burr, Fine Science Tools, Inc.) while under regular artificial cerebrospinal fluid (buffered pH 7.4) perfusion. A 4 mm round #1.5 cover glass (World Precision Instruments, Inc.) was placed over the exposed brain with artificial cerebrospinal fluid filling the void. Gentle pressure was applied to the cover glass while Kwik-Sil adhesive (World Precision Instruments, Inc.) was deposited to bond the glass to the skull. A layer of Vetbond tissue adhesive (3M) was applied over the Kwik-Sil to create a sterile, air-tight seal around the craniotomy and to allow for the restoration of intracranial pressure. C&B Metabond (Parkell, Inc.) was then used to cement a custom titanium head-plate centered around the cranial window for head-constrained awake measurements. The subjects were allowed to recover from surgery and monitored for cranial window integrity and normal behavior for at least four weeks prior to imaging. They were then exposed to head fixation during 20-30 minute sessions over 3-5 days until habituated to locomotion on a linear treadmill awake imaging system [34].

Mice were housed 1-4 per cage in a conventional vivarium maintained at 20^◦^C on 12-hour light-to-dark cycles (07:00 to 19:00). All subjects received standard cage supplements (e.g. cage enclosures, nesting material, and wooden toys) with water and food *ad libitum*. All animal protocols were designed in accordance with the National Institutes of Health Guide for the Care and Use of Laboratory Animals in compliance with ARRIVE guidelines and were approved by the Institutional Animal Care and Use Committee at The University of Texas at Austin.

### 2.3. Awake-to-Anesthetized Blood Flow Measurements

Cranial window implanted mice were head constrained on the awake imaging system and allowed to acclimate for 5-10 minutes. The awake hemodynamic baseline was then defined prior to continuous imaging by acquiring 50 MESI frames, averaging the speckle contrast across each exposure time, and computing the corresponding *ICT* frame using Eq. (1). A custom 3D-printed inhalation nose-cone connected to the anesthesia vaporizer and scavenger was then placed several millimeters away from the subject and used to administer medical air (0.5 L/min) with no isoflurane (0%). Continuous MESI was initiated (*t* = 0) and used to monitor the awake subject for 10 minutes before increasing isoflurane to 2.0% to induce anesthesia. After one minute of exposure (*t* = 11 minutes), the nose-cone was fully positioned over the subject to ensure targeted delivery of isoflurane. After two minutes of exposure (*t* = 12 minutes), a feedback heating pad (55-7030, Harvard Apparatus, Inc.) was placed beneath the subject to maintain a 37 °C body temperature. The now anesthetized subject was continuously monitored with MESI until *t* = 30 minutes, at which point the anesthetized hemodynamic state was defined by acquiring and averaging an additional 50 MESI frames. The subject was then removed from the imaging system and allowed to recover from anesthesia on a heating pad before being transferred back into its cage. Each subject underwent three imaging sessions with at least one full day of recovery between each session.

One animal (Subject 1) underwent an extended routine where the awake and anesthetized states were each imaged for one hour in order to capture the longer-term hemodynamics. Because of the volume of data being recorded, the acquisition was briefly paused prior to the induction of anesthesia to offload data from the solid-state drive used for writing data, which resulted in a short period of missing imagery. The timing of the anesthesia induction was identical to the protocol described above with the awake and anesthetized states defined at *t* = 0 and *t* = 120 minutes, respectively. The four repeated imaging sessions for this subject were spaced at least one week apart to minimize any complications from extended anesthesia exposure.

### 2.4. Data Processing

All data processing was performed using MATLAB (R2021a, MathWorks, Inc.). While head fixation can greatly minimize locomotion-associated brain movement [35], it does not completely eliminate motion artifacts. The induction of general anesthesia also causes major changes in posture that can result in lateral displacements of the brain relative to the skull. In order to account for these shifts, the open source image registration toolbox Elastix [36] was used to rigidly align all speckle contrast frames to the baseline awake imagery prior to fitting Eq. (1). The resulting aligned *ICT* frames were then smoothed temporally with a central moving average filter (*k* = 5) and rendered to video at 20 frames-per-second (8× speed). *ICT* timecourses spanning the entire imaging session were then extracted from the aligned data from both vascular and parenchymal regions of interest (ROIs).

#### 2.4.1. Calculating Vascular Flux

In order to calculate the vascular flux [32], the diameter of vessels (*d*_*vessel*_) were estimated using the 5 ms exposure speckle contrast frames. Cross-sectional profiles were extracted across each vessel of interest at every timepoint and fitted to a Gaussian function in order to calculate the full-width at half-maximum (FWHM) of the distribution over time. Because LSCI is most sensitive to moving scatterers, i.e. erythrocytes, this method of estimating *d*_*vessel*_ is a measure of the inner diameter of the vessel. The vascular flux was then computed by scaling the *ICT* by *d*_*vessel*_ at each timepoint. The percent change in vessel diameter and vascular flux were calculated using the average of the entire awake state for the baseline.

#### 2.4.2. Calculating Blood Flow Rise Time

To examine the dynamics of isoflurane-induced flow changes, the rise time in measured blood flow was calculated. Rise time was defined as the time elapsed following the start of anesthesia for flow values to increase by 50% relative to the awake baseline. A spatially-resolved blood flow rise time map based on *ICT* was generated for Subject 1 by performing this calculation pixel-wise across the entire field-of-view. Analogous rise time calculations were performed on the vascular flux and parenchymal tissue perfusion metrics across all subjects and imaging sessions using the previously extracted ROI timecourses.

#### 2.4.3. Statistical Analysis

All statistical analyses were also performed using MATLAB. The differences between arterial, venous, and parenchymal measurements were evaluated with one-way ANOVAs using the Bonferroni method for multiple comparisons. *p* ≤ 0.05 was considered statistically significant.

## 3. Results

The systematic changes caused by general anesthesia with isoflurane in Subject 1 are shown in Fig. 1b and Video 1, which highlights the transition from the awake to the anesthetized state. The visible decrease in speckle contrast and corresponding increase in *ICT* are representative of an increase in CBF caused by the vasodilatory effects of isoflurane. Dilation is seen across much of the surface vasculature, in particular the large central vein and numerous smaller vessels that have become more prominent in the anesthetized imagery as blood flow increased. These large alterations to the vasculature complicated rigid image registration and precluded the generation of relative flow change imagery.

The temporal dynamics of the awake-to-anesthetized transition can be seen in Fig. 2, where the change in blood flow within two veins (R1 and R2) and one parenchyma region (R3) were analyzed during the extended two-hour imaging session in Subject 1. The cross sections used to estimate *d*_*vessel*_ are depicted in Fig. 2a with the resulting changes in vessel diameter shown in Fig. 2b. The width of the two vessels remained steady throughout the entirety of the awake measurements but increased under anesthesia, with the larger vessel (R1) growing by over 50% after an hour of exposure. This vasodilation was also accompanied by an increase in blood flow as measured by the percent change in vascular flux and parenchymal tissue perfusion as shown in Fig. 2c. The vascular flux in R1 and R2 increased by 270% and 144%, respectively, while the tissue perfusion in R3 grew by 141%. The large deviation shortly after *t* = 80 minutes was caused by the subject briefly experiencing breathing difficulties that required the anesthesia be reduced to 1.5% isoflurane for several minutes. The breathing abnormality and change in anesthesia dosage had minimal effect on the vessel size. Repeated measurements in the subject (Figs. 2d-f) illustrate the stability of day-to-day MESI estimates of blood flow as well as the consistency of the hemodynamic response to isoflurane anesthesia.

**Figure 2:**
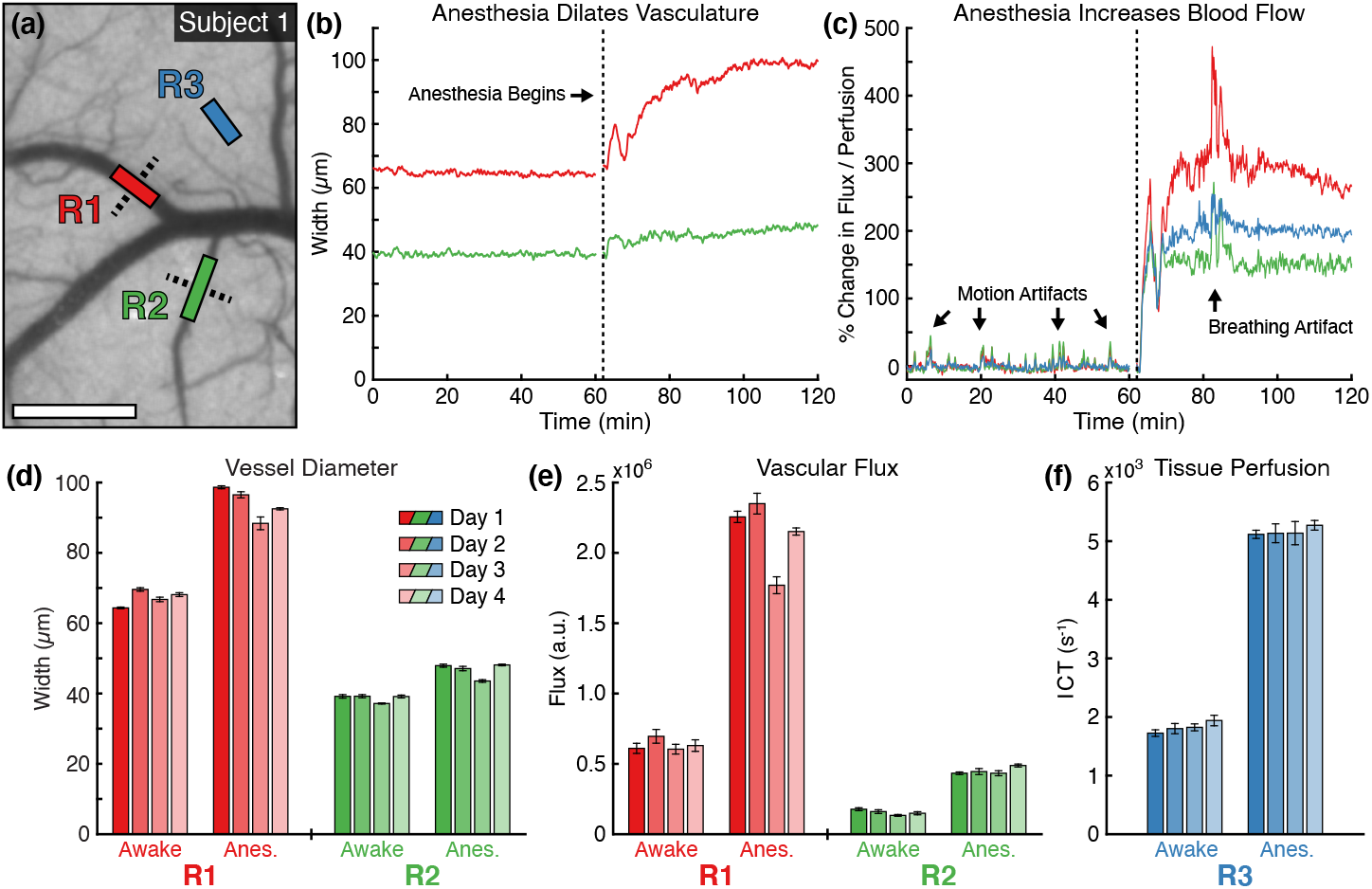
(a) Awake speckle contrast image (5 ms) from Subject 1 overlaid with vascular (R1 and R2) and parenchyma (R3) regions used during flow analysis (Scale bar = 500 *µ*m). Dashed lines indicate the cross-sectional profiles used to calculate vessel width. Timecourses of (b) vessel diameter and (c) percent change in blood flow within the regions over the extended two-hour imaging session. The vertical dashed line indicates the beginning of anesthesia induction. The percent change in vascular flux and tissue perfusion in (c) were computed using the average of the entire awake state for the baseline. Repeated measurements of (d) vessel diameter, (e) vascular flux, and (f) parenchyma tissue perfusion during awake and anesthetized states in Subject 1 across four imaging sessions (mean ± sd). Statistics computed across the 50 MESI frames acquired in each state.

The percent changes in vessel diameter and blood flow between the awake and anesthetized states were aggregated across all subjects and imaging sessions as shown in Fig. 3. The anesthetized measurements for Subject 1, which underwent the extending imaging sessions, were taken at *t* ≈ 80 minutes to match the timing of the other subjects (20 minutes after the start of anesthesia). A total of 108 measurements were included in the analysis (33 arteries, 29 veins, 46 parenchyma regions). Similar to the results seen in Fig. 2, the vasodilatory effects of isoflurane induced large changes in the cortical hemodynamics. On average, vessel width increased by 14.1%, with arteries increasing by 15.0% and veins increasing by 13.0% (Fig. 3a). There was no significant difference in the percent change in width between arteries and veins (*p* = 0.61). Vascular flux increased, on average, by 96.0%, with arteries increasing by 81.7% and veins increasing by 112.2% (Fig. 3b). Parenchymal tissue perfusion increased by 84.7%. While the ANOVA was significant (*p* = 0.041), the multiple pairwise comparisons between arteries, veins, and parenchyma regions were all non-significant (*p >* 0.05).

**Figure 3:**
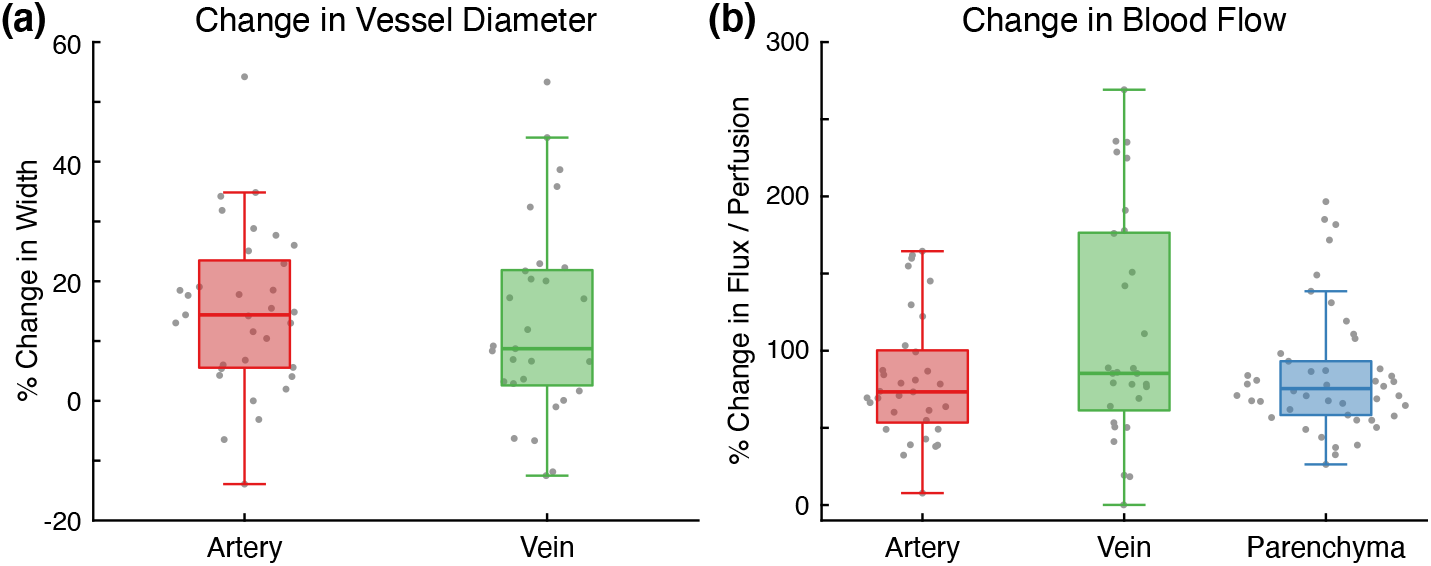
(a) Percent change in vessel diameter for arteries and veins between the awake and anesthetized states (*p >* 0.05). (b) Percent change in arterial and venous vascular flux and parenchymal tissue perfusion between the awake and anesthetized states (*p >* 0.05). Measurements in both charts aggregated across all subjects and all imaging sessions.

The dynamics of the isoflurane-induced flow changes in Subject 1 can be seen in the blood flow rise time map in Fig. 4a. *ICT* values across the entire field-of-view increased by at least 50% in just over one minute following the start of anesthesia. The vasculature, including both arteries and veins, appears to change flow faster than parenchyma regions. However, upon aggregating ROI measurements across all subjects and imaging sessions (Fig. 4b), arteries have significantly faster rise times (40.5 seconds) compared to veins (50.7 seconds, *p* = 3.7 × 10^−6^) or parenchyma regions (46.1 seconds, *p* = 0.006). Parenchymal regions also have significantly faster rise times than veins (*p* = 0.043).

**Figure 4:**
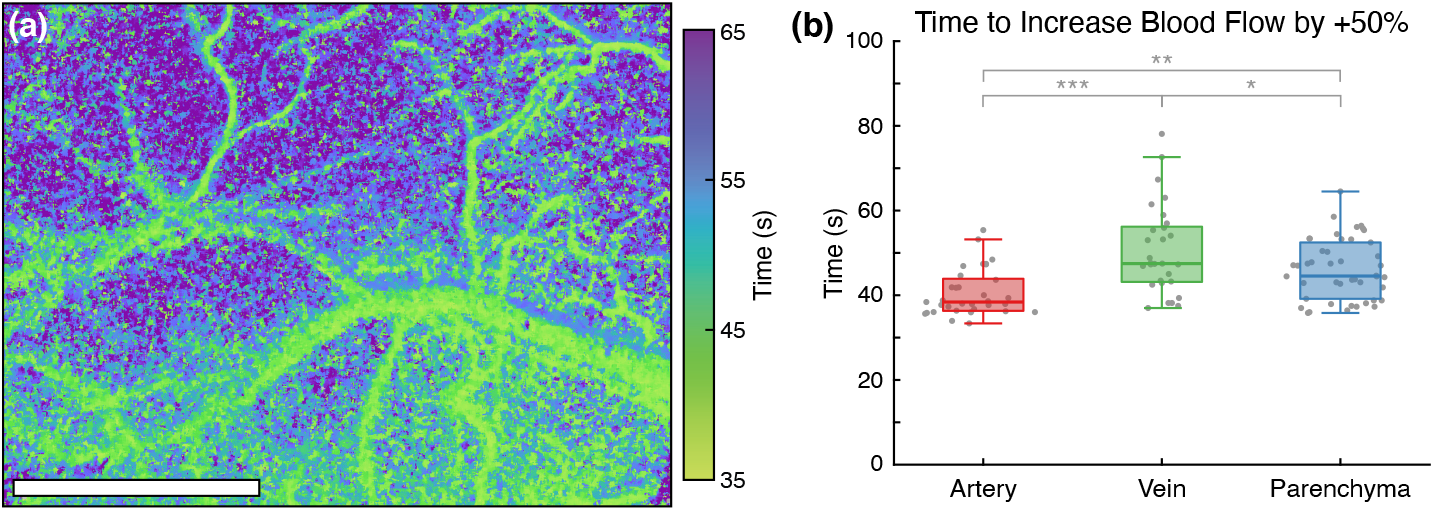
(a) Blood flow rise time map depicting the time elapsed following the start of anesthesia for *ICT* to increase by 50% relative to the awake baseline in Subject 1 (Scale bar *×* = 1 mm). (b) Time for arterial and venous vascular flux and parenchymal tissue perfusion to increase by 50% relative to the awake baseline (\*\*\**p* = 3.7 × 10^*−*6^, \*\**p* = 0.006, \**p* = 0.043). Measurements aggregated across all subjects and all imaging sessions.

## 4. Discussion

The systemic effects of isoflurane undermine the reliability of neurophysiological and behavioral studies because anesthesia is a fundamentally unnatural physiological state [17]. Imaging blood flow on the cortical surface with MESI during the induction of anesthesia directly visualizes and quantifies these large hemodynamic changes. On average, the vasodilation caused by isoflurane increased vessel diameter by 14.1% across all trials, with arteries increasing by 15.0% and veins increasing by 13.0% (Fig. 3a). Direct microscope measurements in fentanyl- and nitrous oxide-anesthetized rats documented a 17% increase in arteriolar diameter and 6% increase in venule diameter after the topical application of isoflurane [8]. Similar measurements in pentobarbital-anesthetized dogs observed 10-28% increases in the diameter of small arterioles following the inhalation of isoflurane [9]. Both studies found that isoflurane dilated arterioles in a concentration-dependent manner. More recent measurements in mice using optical coherence tomography (OCT) angiography found that isoflurane increased artery diameter by 12-55% and vein diameter by 14-22%, depending on vessel branch order [37].

Much larger changes were measured in surface vessel blood flow, with vascular flux increasing on average by 96.0% across all trials (Fig. 3b). This suggests that increases in blood flow velocity rather than vasodilation are largely responsible for the measured change in flow. Because few imaging modalities are capable of measuring the dynamics of blood flow with sufficient spatial resolution to distinguish individual vessels, there has been limited work on this topic in awake animals. One study using laser Doppler flowmetry found only non-significant 18% increases in both CBF and red blood cell (RBC) velocity in the barrel cortex after mice were anesthetized with isoflurane [6]. However, because large blood vessels were avoided, the laser Doppler measurements only sampled smaller subsurface vasculature that may exhibit different responses to isoflurane than the larger surface vasculature imaged in this study. The vascular flux metric calculated from the speckle correlation time is also not analogous to either the CBF or RBC velocity measured by laser Doppler flowmetry, so the results are not directly comparable [32]. Another more recent study using Doppler OCT measured a 55% increase in volumetric blood flow in mice under isoflurane [37]. Unlike the results presented above, this change was largely driven by vasodilation rather than an increase in blood flow velocity.

Within the parenchyma, tissue perfusion increased on average by 84.7% across all trials after isoflurane exposure (Fig. 3b). The narrower distribution of values compared to the vascular measurements is likely a byproduct of the more homogenous structure of the unresolvable capillaries within the parenchyma. fMRI measurements in isoflurane-anesthetized rats have reported ∼50% increases in global CBF with the cerebral cortex experiencing the smallest regional increase (20%) compared to the awake state [38]. However, these numbers are not directly comparable because fMRI samples a much larger volume of the brain than MESI, which has both a smaller field-of-view and only penetrates several hundred microns into the cortex.

While LSCI has been utilized extensively with awake imaging [39, 16, 14, 20, 40], this is one of the first uses of MESI in an awake animal model [34]. Traditional single-exposure LSCI would likely have underestimated the magnitude of the flow changes caused by isoflurane because individual exposure times are only sensitive to a narrow range of flow rates [21]. Failing to account for the increase in multiple dynamic scattering events caused by vasodilation would also have resulted in an underestimation of the flow changes in the surface vasculature. Using the raw *ICT* value instead of the diameter-scaled vascular flux metric in Fig. 2c would have measured only a 141% increase in the large vessel instead of 270%. The repeatability of the day-to-day measurements shown in Figs. 2d-f further demonstrate the robustness of MESI as a chronic blood flow imaging technique [23], even when observing a fully-conscious, moving animal.

The generation of the blood flow rise time map (Fig. 4a) highlights the continuous wide-field imaging capabilities of MESI. Other blood flow imaging techniques such as laser Doppler flowmetry, OCT, or fMRI would be restricted by either spatial or temporal resolution from performing such an analysis. The spatial heterogeneity in the mapping reveals differing dynamics between the vasculature and parenchyma as confirmed with the aggregated ROI analysis in Fig. 4b. While image registration was used to spatially align the *ICT* frames, the isoflurane-induced vasodilation complicated this process and resulted in imperfect alignment. The effects of this are most pronounced along the periphery of vasculature where the vasodilatory changes were the most prominent.

The effects of isoflurane fundamentally undermine the imaging of cortical hemodynamics in the normal physiological state. Blood flow imaging techniques such as LSCI, laser Doppler flowmetry, and fMRI require greater sensitivity because of the suppression of the hemodynamic response [6] and measure a delayed neurovascular coupling [7]. Anatomical imaging with two-photon microscopy only captures heavily-dilated vasculature unrepresentative of the awake resting state [19]. The consistency and reliability of all imaging techniques can suffer from the effects of prolonged isoflurane exposure, which can cause continual vasodilation as seen in Fig. 2b. This could cause extended imaging sessions to document disparate anatomies and physiologies between the beginning and conclusion of an experiment.

### 4.1. Limitations

The impact of repeated exposure to isoflurane was not examined in this study. Previous work has found that it can impair synaptic plasticity in the basolateral amygdala [41] and cause persistent motor deficits via structural changes to the corpus callosum [42]. The one day interval between imaging sessions may not have been sufficient to completely avoid these effects despite only 20 minutes of isoflurane exposure. Because the experiment imaged the induction of general anesthesia from the awake state, tracheal intubation for mechanical ventilation was not an option. This may have resulted in breathing variability that impacted the systemic hemodynamics and complicated comparisons with other studies that had mechanical control of ventilation. Since varying the isoflurane dosage was the only method of regulating breathing stability, the concentration dependent effects of isoflurane may have also influenced the results [43, 44].

While image registration can help maintain the spatial alignment of data across an experiment, it is unable to decouple the effects of animal motion from the underlying blood flow. Even with the head fully restrained, walking and grooming both caused subtle brain movements that manifested as large changes in speckle contrast, as seen by the abrupt spikes during the awake section of Fig. 2c. While these fluctuations may represent real hemodynamic responses, it is difficult to isolate them from broader animal motion. A simple solution might be the addition of an external sensor to the treadmill to exclude timepoints when the animal is actively moving [35]. However, a more refined awake imaging system that further minimizes brain movement would likely be necessary to directly probe these phenomena with LSCI or MESI.

## 5. Conclusion

We have used MESI to continuously image CBF during the induction of general anesthesia with isoflurane in head-restrained mice. The vasodilatory effects of isoflurane caused rapid and large anatomical and physiological changes that were drastically different from the awake state. These increases in vessel size and blood flow were repeatable across multiple imaging sessions and subjects. We also documented disparate response times to the induction of isoflurane anesthesia between arteries, veins, and parenchyma regions. This study demonstrated that MESI can be readily used for chronic awake imaging and allows for direct day-to-day comparisons of blood flow. These results provide further evidence that the anesthetized state is unrepresentative of normal physiology and that neurophysiological and behavioral experiments would benefit immensely from transitioning to un-anesthetized awake animal models.

## Acknowledgements

This study was supported by the National Institutes of Health (Nos. EB011556, HL140153, NS108484, NS109361, and T32EB007507).

## Declaration of Interest

A.K.D. holds equity in Dynamic Light, Inc. The authors declare no other conflicts of interest.

### CRediT Statement

**Colin T. Sullender:** Conceptualization, Methodology, Software, Validation, Investigation, Data Curation, Formal Analysis, Writing - Original Draft, and Visualization. **Lisa M. Richards:** Conceptualization, Methodology, Investigation, Data Curation, and Writing - Review & Editing. **Fei He:** Investigation, Resources, and Writing - Review & Editing. **Lan Luan:** Writing – Review & Editing, Supervision, and Funding Acquisition. **Andrew K. Dunn:** Conceptualization, Writing - Review & Editing, Supervision, Funding Acquisition, and Project Administration.

